# A deep learning approach to pattern recognition for short DNA sequences

**DOI:** 10.1101/353474

**Authors:** Akosua Busia, George E. Dahl, Clara Fannjiang, David H. Alexander, Elizabeth Dorfman, Ryan Poplin, Cory Y. McLean, Pi-Chuan Chang, Mark DePristo

## Abstract

**Motivation:** Inferring properties of biological sequences--such as determining the species-of-origin of a DNA sequence or the function of an amino-acid sequence--is a core task in many bioinformatics applications. These tasks are often solved using string-matching to map query sequences to labeled database sequences or via Hidden Markov Model-like pattern matching. In the current work we describe and assess a deep learning approach which trains a deep neural network (DNN) to predict database-derived labels directly from query sequences.

**Results:** We demonstrate this DNN performs at state-of-the-art or above levels on a difficult, practically important problem: predicting species-of-origin from short reads of 16S ribosomal DNA. When trained on 16S sequences of over 13,000 distinct species, our DNN achieves read-level species classification accuracy within 2.0% of perfect memorization of training data, and produces more accurate genus-level assignments for reads from held-out species than *k*-mer, alignment, and taxonomic binning baselines. Moreover, our models exhibit greater robustness than these existing approaches to increasing noise in the query sequences. Finally, we show that these DNNs perform well on experimental 16S mock community dataset. Overall, our results constitute a first step towards our long-term goal of developing a general-purpose deep learning approach to predicting meaningful labels from short biological sequences.

**Availability:** TensorFlow training code is available through GitHub (https://github.com/tensorflow/models/tree/master/research). Data in TensorFlow TFRecord format is available on Google Cloud Storage (gs://brain-genomics-public/research/seq2species/).

**Supplementary information:** Supplementary data are available in a separate document.

## 1 Introduction

Many important problems in bioinformatics--including taxonomic profiling, protein family assignment, and detection of regulatory elements--require a method of accurately inferring meaningful labels from short sequences, which we call the sequence-labeling problem. A common approach to labeling these query sequences is to build a database of reference sequences for the property of interest, map the unknown queries to the database using *k*-mer matching or pairwise-sequence alignment, and finally use the mappings and their scores to assign labels. For example, many top-performing taxonomic binning and profiling tools use an initial *k-*mer mapping or local alignment of each read and then integrate these per-read mappings into higher-level taxonomic mixture estimates.^5, 32-35^ Hidden Markov Models (HMMs), which are widely used for annotating genes and regulatory elements, represent an alternative approach wherein one builds a statistical model from the relevant database and then uses this model to assign labels to queries.^38-40,49^ Here, we aim to assess how valuable deep learning might be in approaching these types of sequence-labeling problems using taxonomic classification of short reads of 16S ribosomal DNA as an initial benchmarking problem due to its appropriate level of difficulty and significant practical applications.

The flexibility of deep neural networks (DNNs) and their ability to learn complex patterns make them a promising choice for solving the general sequence-labeling problem. String-matching approaches work well, but are often unable to accurately capture the full complexity of the sequence/label relationship--for example due to loss of positional information in *k*-mer methods or oversimplified affine gap penalties^41^ used in alignment tools like the Basic Local Alignment Search Tool (BLAST)^27^ or Burrows-Wheeler Aligner (BWA).^25,26^ Recent work using DNNs to learn regulatory elements patterns^42^ provide further evidence that neural networks are better at modeling complex dependencies between sequence positions than methods like HMMs.^38-40,49^ Moreover, using DNNs for sequence-labeling could provide significant performance advantages by leveraging hardware accelerators. We, therefore, design a new DNN architecture to solve the sequence-labeling problem, and explore its behavior on an important example problem to gain initial insights into its behavior and potential utility to the bioinformatics community.

Many sequence-labeling problems in bioinformatics could be used to benchmark our proposed DNN; here we explore that of assigning species-level taxonomy to short reads of 16S ribosomal DNA. This task is inherently challenging due to the presence of closely related organisms and conserved regions, and its difficulty can be easily modulated by holding out information based on phylogeny. Moreover, this problem is both well-characterized--having been used in microbial phylogeny for over forty years^1,2^--and practically important--evidenced by growing public repositories of 16S sequencing data^3,4^ and the recent Critical Assessment of Metagenomic Interpretation (CAMI).^7^ CAMI’s top-performing taxonomic binning and profiling tools use initial *k*-mer matching (PhyloPythiaS+^32^and Kraken^33^) and local alignment (taxator-tk,^34^ MEGAN,^5^ MetaPhlAn,^6^ and MetaPhyler^35^) steps. Other existing approaches to read-level taxonomic labeling use *k*-mers via naive Bayes, probabilistic topic models, and Markov models, but tend not to outperform BLAST for typical sequence lengths.^8,9,31,36^Early studies using neural networks for taxonomic classification from DNA barcodes provide initial support for a deep learning approach, but lack sufficient data and experimental validation to provide meaningful guidance on whether a modern DNN solution has any practical advantages over existing tools.^10,11^

Despite the recent increase in successful applications of deep learning to genomics,^46-48^ to our knowledge no previous work demonstrates that DNNs can classify genomic sequencing data of realistic scales at fine taxonomic resolution, or provides an in-depth comparison to existing sequence-matching and read-level taxonomic binning techniques. We show that our proposed neural networks architecture scales successfully to more than 13 thousand distinct species, and moreover that it can accurately analyze both synthetic and experimental read data. We find that these learned, discriminative classifiers outperform *k*-mer based classifiers and achieve performance which is comparable to that of traditional alignment tools, while also being more robust to both increasing noise and decreasing read length. We show our DNNs are also more robust than Kraken, a popular taxonomic binning tool. We use these results to understand the conditions under which this deep learning solution is most advantageous, and conclude by discussing the generality of our approach.

## 2 Methods

### Data

We used 19,851 16S ribosomal RNA reference sequences from the NCBI RefSeq Targeted Loci Project^20^ (BioProjects 33175 and 33317, downloaded 2017-11-27). For *L =* {25, 50, 100, 150, 200}, we constructed a set of synthetic reads of length *L* (denoted NCBI_L_) by extracting all subsequences of *L* base pairs from each reference sequence. Each read is paired with superkingdom, phylum, class, order, family, genus, and species labels from NCBI Taxonomy Browser.^21^ The data represents 13,838 species from 2,768 genera (Appendix 1). For experiments on held-out species, we split NCBI_L_ into training and validation subsets containing 12,598 and 1,230 species, respectively. For model selection, we split NCBI_L_ into three subsets (Appendix 3).

To evaluate our trained classifiers, we used 16S mock community sequencing data from Nelson et al.,^22^ Schirmer et al.,^29^ and D’Amore et al.^23^ (PRJEB4688 and PRJEB6244 in the European Nucleotide Archive^24^). We used six Illumina MiSeq replicates (ERR348713-5 and ERR619081-3) of the 20-organism mock community (PRJEB4688) and all 51 runs from the 59-organism community (PRJEB6244). Appendix 7 details the contents of each replicate.

### Model Architecture

Each canonical base (A, C, T, G) or IUPAC ambiguity code in a given input was encoded as an appropriate four-dimensional probability distribution over the four canonical bases (Extended Data Figure 1). Similarly, we one-hot encoded species labels as a 13,838-dimensional vector.

**Figure 1:**
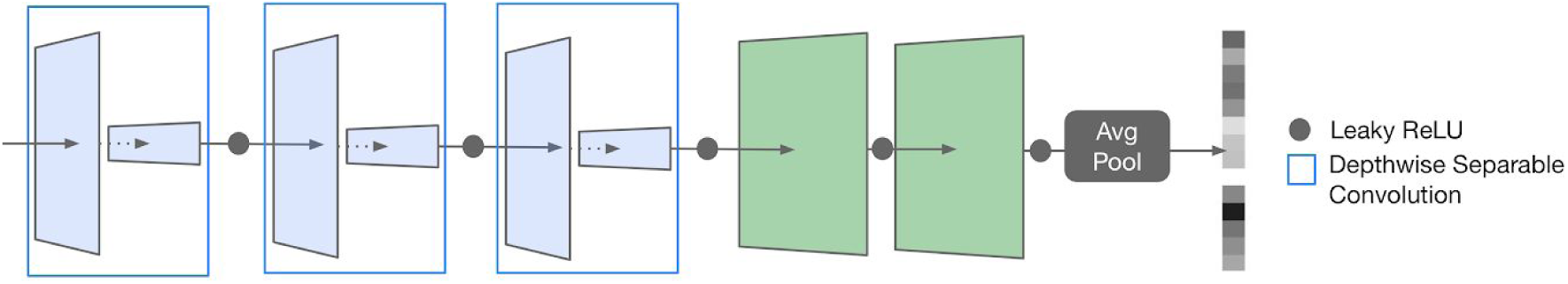
Overview of the proposed neural network architecture. The neural network architecture used in this study consists of three depthwise separable convolutional filters followed by two or three fully-connected layers (in green), which are tiled as needed along the length of the input and combined via an average pooling layer prior to the softmax output layer.

We used a model architecture comprised of three layers of depthwise separable convolutions (sequential spatial and pointwise convolution operations), two to three fully-connected layers, a pooling layer, and a softmax output layer that produces a probability distribution over possible species labels. We apply dropout regularization^30^ and leaky rectified-linear activation^17, 18^ to each convolutional and fully-connected layer (Appendix 2). When inputs are longer than the width of the fully-connected layers, these layers are tiled as needed and the pooling layer combines the intermediate outputs before the softmax output layer. We found that average pooling worked best for this application (Appendix 4). Appendix 3 describes model selection and gives the best configuration we found for each minimum read length. To compute probability distributions over higher taxonomic labels, we marginalized the predicted species-level distributions as follows:

Given an input read *x*, fixed model *DNN*, and higher-order taxon of interest:

1. Let *DNN*(*x*)_*i*_ denote the predicted probability assigned by the model to the *i*^th^ species label, and take *I*_*j*_={*i* : *i*^th^ species is contained in *j*^th^ higher-order taxon}
2. Return higher-order probabilities *P*(*x*) computed as 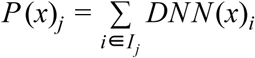

### Training and Implementation

We implemented our neural networks using the open-source software library TensorFlow.^16^ Starting with randomly initialized parameters (Appendix 2), we used a random mini-batch of 500 read and species label pairs on each iteration to update model parameters to minimize cross entropy loss between the true and predicted species labels (using TensorFlow’s implementation of the ADAM optimizer^19^).

We also explored classification performance as a function of noise rate. When we injected random base-flipping noise into input sequences, we mutated each base *b* with fixed probability *r* according to the following rule:

If *b* is a canonical base:

Flip *b* to one of the other three canonical bases with equal probability.

Otherwise:

Flip *b* to one of the four canonical bases with equal probability.

We trained models with five different noise rates *r* = {0%, 2%, 4%, 8%, 16%} and evaluated on data with injected noise rates of *r* = {0%, 0.5%, 1%, 2%, 5%, 10%}.

### Baselines

We computed the Bayes optimal accuracy for a fixed set *T* of training examples as follows:

1. Partition *T* into subgroups *T* _1_, *T* _2_,…, *T*_*N*_ of (read, species label) pairs, where *N* is the number of distinct read sequences, and all training examples in *T* which share the same read sequence are contained in the same partition *T*_*i*_
2. Define *count*_*i*_(*s*) as the number of times the species label *s* appears in the subgroup *T*_*i*_ and compute 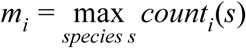 for *i* ∈ {1, 2,…, *N*}
3. Let |*T* | be the total number of reads in *T* and take 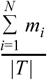 as the final accuracy We repeat this process for each higher taxonomic rank.

For our alignment baselines, we used BLAST^27,28^ and BWA^25,26^ to align queries against the original NCBI reference sequences and computed the accuracy of randomly guessing a label for each read from the set of labels for all references with equally good mappings. For BWA, we consider all references with edit distances as least as good as the primary mapping, and for BLAST we use the bit score (Appendix 6).

Our naive Bayes classifier (based on the RDP Classifier^8^) uses 8-mer matches of queries reads to the original references sequences. We transformed input sequences into vectors indicating the presence or absence of each possible 8-mer subsequence. For each such 8-mer, the prior probability used for its presence and genus-specific conditional probability were computed as in Wang et al.^8^ We handled IUPAC ambiguity codes in the 8-mer vector representation by assigning each possible DNA 8-mer a fractional presence (Appendix 6).

We also compared the sensitivity, positive predictive value (PPV), and F1 scores of our DNNs to Kraken2^33^(https://github.com/DerrickWood/kraken2, downloaded 2018-09-28) a popular taxonomic binning tool that emits read-level predictions. We generated a kraken2 database in exact correspondence with the taxonomy of organisms in our NCBI training data (Appendix 6), and ran kraken2 on FASTQ files of query sequences using the following command:

~~~
$ kraken2 --db kraken2_database --threads 48 kraken.fastq > output.kraken
~~~

kraken2 emits whether each read was classified (C) or unclassified (U), and gives a predicted taxonomic rank and entity for each classified read. All ranks above the emitted rank are marked with the entity implied by traversing up the taxonomy from the predicted rank, while all ranks below the predicted rank are marked as no-call. We calculated counts of true positives and false positives, along with derived sensitivity, specificity, PPV, false discovery rate, and F1 values at each rank by comparing to the true taxonomic rank and entity while respecting no-calls (i.e. sensitivity includes no-calls as false negatives while specificity ignores no-calls).

### Mock Community Evaluation

Base-flipping noise does not capture all the interesting aspects of real sequencing errors;^37^ we thus also applied our DNNs to experimental reads with real error profiles. We used the following iterative method to obtain mixture estimates from read-level predictions:

1. Using a fixed model, *DNN*, initialize a matrix of read-level probabilities *P* so that for each row row *i, P _i_* = *DNN* (*x*_*i*_) where *x*_*i*_ is the *i*^th^ read in the set of raw sequencing reads.
2. Initialize a vector of mixture estimates *E* as the uniform distribution over possible species labels.
3. For each iteration:
  a. Compute a matrix *T* where *T _i_* = (*E* °*P _i_*) / (*E · P _i_*) where ° is element-wise multiplication
  b. Update 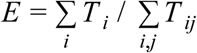

We used 10,000 reads and 75 iterations for the smaller mock community, and 20,000 reads and 150 iterations for the larger. We took the list of species with estimated mixture fractions over 0.001 in the final *E* as the predicted set of community members. We repeated this procedure to estimate genus-level composition by marginalizing the model’s predictions to the genus level, and initializing *E* as a uniform distribution of the appropriate size.

## 3 Results

### Training Performance

For each of five minimum read lengths *L* = {25, 50, 100, 150, 200}, we explored whether a neural network, DNN_L_, of the structure in Figure 1 can learn to predict accurate read-level labels. While our architecture’s pooling layer allows the models to tolerate variation in read length, we found it informative to train multiple models optimized for different lengths (Figure 2b). We trained each DNN_L_ independently on the synthetic reads in NCBI_L_.

**Figure 2:**
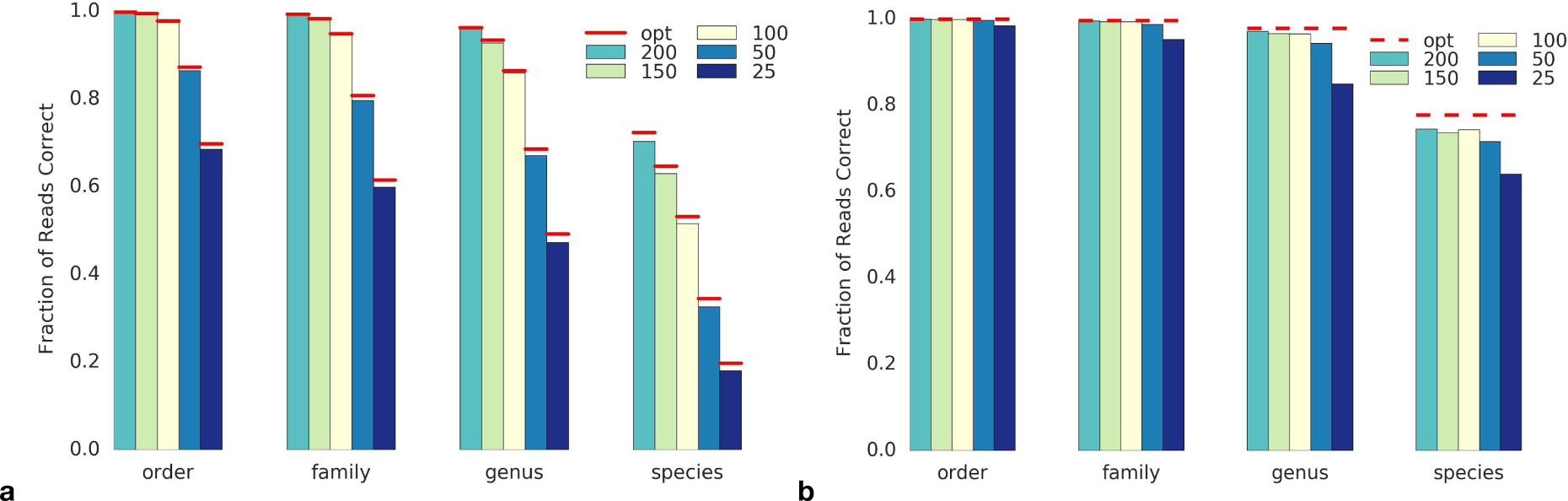
Neural network read-level accuracy relative to Bayes optimal solution. **(a)** The results of each neural network DNN_200_, DNN_150_, DNN_100_, DNN_50_, and DNN_25_ on its training set (NCBI_200_, NCBI_150_, NCBI_100_, NCBI_50_, and NCBI_25_, respectively) on the order, family, genus, and species predictions tasks compared to the Bayes optimal accuracy rate (‘opt’). **(b)** The same as **(a)** when all methods are evaluated on reads from NCBI_250_.

We compare to the Bayes optimal classifier, which gives the maximum achievable accuracy on the training set for a perfect read mapping. Overall, classification accuracy increased with read length, and each DNN achieved species classification accuracy within 2.0% of the Bayes optimal rate (Figure 2a). Species-level classification accuracies using BWA fell, on average, 4.6% below Bayes optimal. Higher-order predictions obtained by marginalizing the models’ predicted probability distributions over species (Methods) were also highly accurate, coming within 1.0%, 0.6%, and 0.5% of Bayes optimal rates at the genus, family, and order levels, respectively.DNN_25_attained 84.4%, 90.0%, and 99.9% read-level accuracies on class, phylum, and superkingdom classification; the remaining models surpassed 95.7% accuracy on all these tasks.

On NCBI_250_ all the models performed well at higher-order classification tasks, attaining upwards of 98.3% read-level accuracy on order prediction. At the species-level, models trained on longer reads achieved noticeably better performance on NCBI_250_, falling within 4.1% of the Bayes optimal rate, compared to 6.1% and 13.6% for DNN_25_ and DNN_50_, respectively.

### Experiments on Held-out Species

We retrained DNN_25_, DNN_50_, DNN_100_, and DNN_200_ on synthetic reads from a subset of 12,598 of the 13,838 species in the NCBI data, with base-flipping noise randomly injected into inputs at train time. We compared the efficacy of these models at making genus-level predictions for novel species to alignment and naive Bayes baselines (Methods). Figure 4 shows genus-level accuracy for each method on synthetic reads in the validation set, comprised of synthetic reads from 16S references of the 1,240 species held-out from the training set. The first column of Figure 4 shows that genus-level accuracy on errorless validation reads varies significantly across the different methods, with our approach significantly outperforming for every read length except 200 base pairs where the results are roughly comparable to alignment.

**Figure 3:**
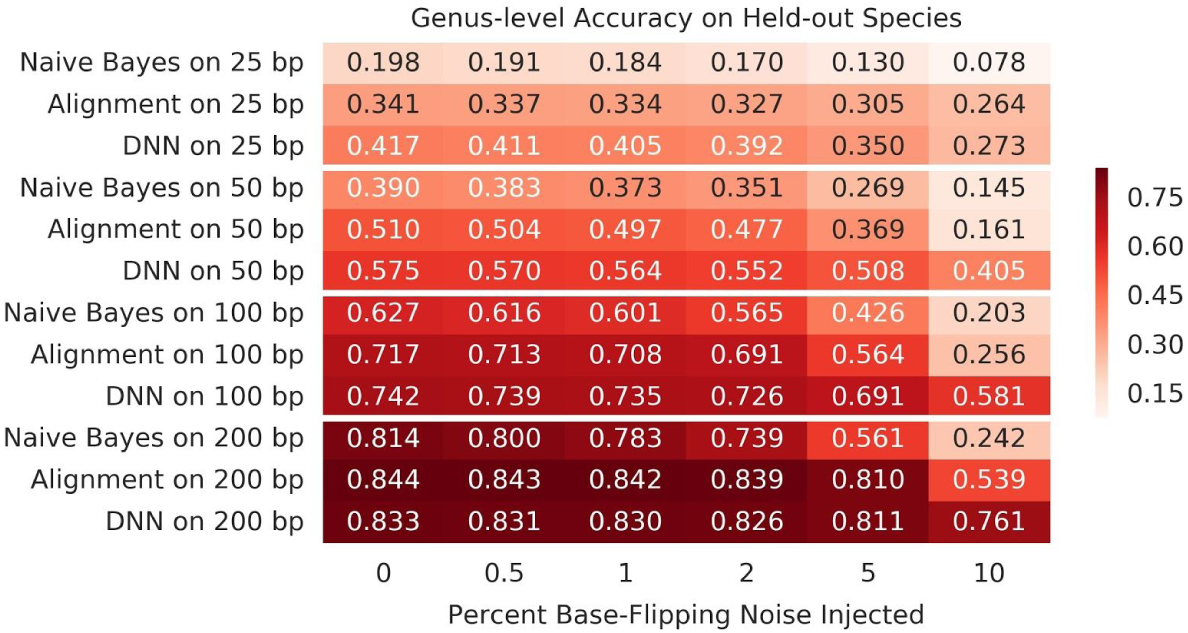
Accuracy and consistency of our deep learning approach on reads from held-out species relative to existing tools. Heatmap showing the read-level accuracy of our deep learning approach (using 8% noise injection during training for 200 base pairs and 4% for all others) relative to naive Bayes and alignment approaches on the task of assigning reads from held-out species to correct genera for several combinations of read length and noise rate. Here, alignment represents the better of our BLAST and BWA baselines, and each cell is labeled with the proportion of reads placed into the correct genus when the corresponding method (*y*-axis) is used to assess held-out reads corrupted by the given noise rate (*x*-axis).

Across the 24 read length and noise rate pairs we tried, our DNNs outperformed the naive Bayes baseline by between 1.9% and 51.9% (Figure 3). In 20 of these 24 settings, our DNNs outperformed alignment baselines; these improvements ranged from 0.1% to 32.5%. For the remaining 4 low-noise experiments on 200 base pair reads, BLAST attained no more than a 1.3% advantage over DNN_200_. Our DNNs performed well even when BWA failed to align a significant portion of the reads; for example, BWA achieved less than 3% read-level genus accuracy on 200 base pair validation reads with 5% or 10% injected base-flipping noise, whereas DNN_200_ achieved 81.1% and 76.1% accuracy, respectively. Thus, in general we found that, compared to *k*-mer and alignment baselines, our proposed deep learning approach tends to assign more accurate genus labels to reads from novel species and is more robust to sequencing errors and the high mutation rate involved in bacteria genomes.

### Kraken Comparison

Figure 4 compares the sensitivity, PPV, and F1 scores of our DNNs to Kraken2^33^ on the validation reads from held-out species. Because our method predicts a genus label for every read, its scores are equivalent across the three performance metrics. In contrast, Kraken emits labels for only a subset of the reads, in general allowing it to attain high PPV at the cost of lower sensitivity. Kraken2’s PPV was highest on validation sets with low noise rates; once the noise rate increased upwards of 1%, the PPV attained by our DNNs was comparable in most cases, and higher for 100 and 200 base pair reads. Sensitivity achieved using Kraken2 was much lower than our DNNs, even when no noise was added to the validation reads. Indeed, our proposed approach achieved high sensitivity and good PPV in general, resulting in much higher F1 scores on genus-level prediction for the held-out species compared to Kraken for every combination of read length and noise rate evaluated in Figure 4. The performance of our deep learning approach was thus more consistent and degraded less with increasing noise.

**Figure 4:**
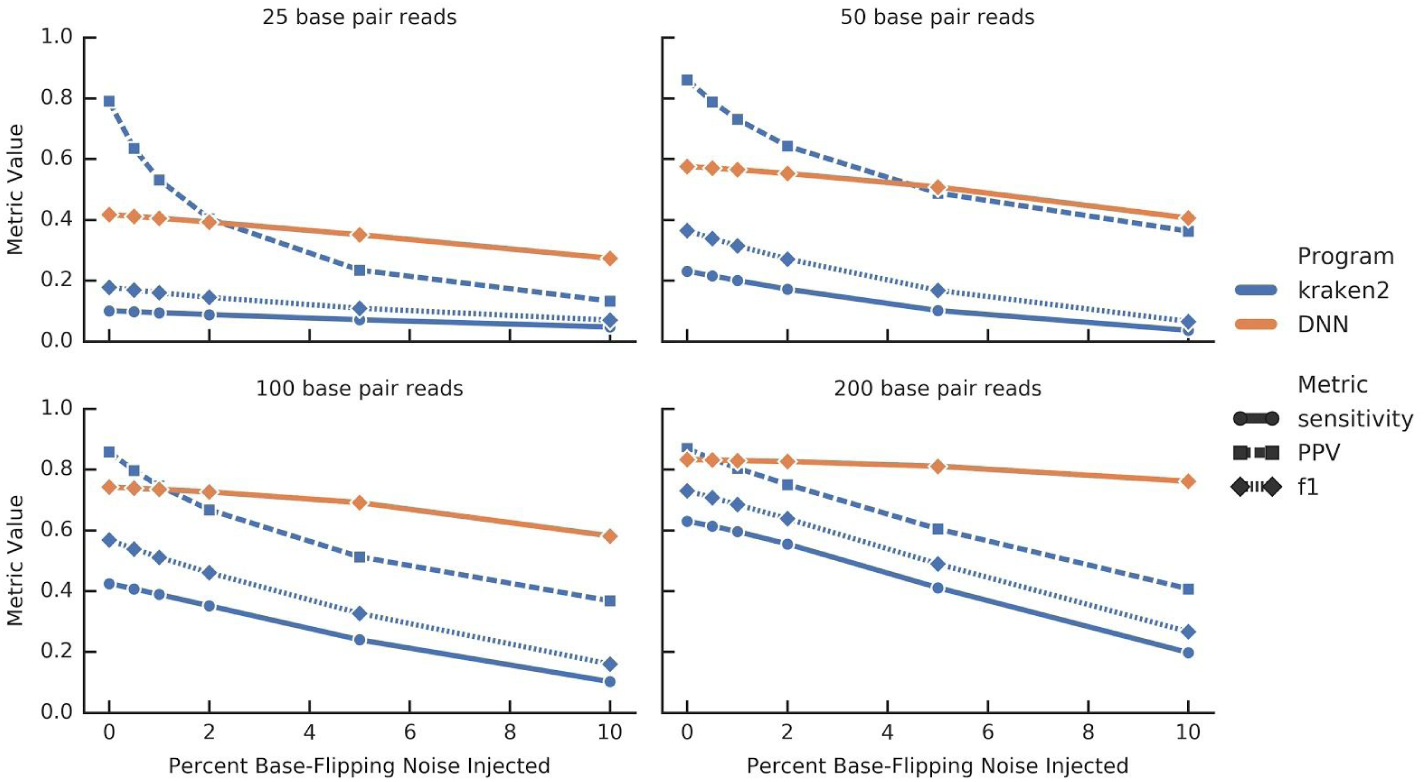
Read-level performance of our deep learning approach relative to Kraken on reads from held-out species. Plots relating the read-level sensitivity, positive predictive value (PPV), and F1 score of kraken2 to the performance our deep learning approach on the task of assigning reads from held-out species to correct genera. Deep learning models, read lengths, and noise rates are consistent with Figure 3. Note that for our DNNs only a single line is drawn in each figure since sensitivity and PPV (and hence F1) are identical.

### Mock Community Evaluation

Using DNN_100_ to analyze 57 sets of empirical mock community sequencing data,^22,23,29^ we demonstrated that our DNNs achieve reasonable performance on real reads generated using a variety of typical next-generation sequencing protocols. Due to the lack of read-level ground-truth labels for the mock community sequencing reads, we introduce a method of integrating over our model’s read-level predictions to produce model-based community reconstructions (Methods). We chose to keep this model-based estimation method as simple as possible so results directly reflect the model’s sequence-labeling performance, but note that integrating the model’s outputs into a more sophisticated analysis pipeline (for instance, inspired by MetaPhlAn^6^ or MetaPhyler^35^) would likely improve profiling results.

For five of six 20-organism mock community replicates, our model-based estimation perfectly reconstructed the list of 17 genera (Figure 5b). At the species level, our method identified, on average, 21.8 distinct species (16.3 truly present and 5.5 false discoveries). For the remaining 51 mock community sequencing datasets, we correctly identified on average 36.0 of 45 genera and 32.6 of 56 species based on the model’s probability assignments. We consistently failed to recover six Archaeal genera: *Nanoarchaeum* (which is not contained in our NCBI dataset), *Methanocaldococcus, Pyrococcus, Sulfolobus, Archaeoglobus*, and *Ignicoccus*. This is likely due, in part, to the “failure of the V1-V9 primers in amplifying the Archaea”^23^and low representation of Archaea in the NCBI dataset. Overall, across all 57 sets of amplicon sequencing data, our model-based estimation method attained mean PPV and sensitivity of 0.657 and 0.610 at the species level, and 0.908 and 0.824 at the genus level. Though trained on a smaller reference database, compared to RDP Classifier results from D’Amore et al., ^23^our approach discovered fewer spurious genera (4.1 compared to 55.6) without a large decrease in the number of accurately recovered genera (36.0 versus 39.3).

**Figure 5:**
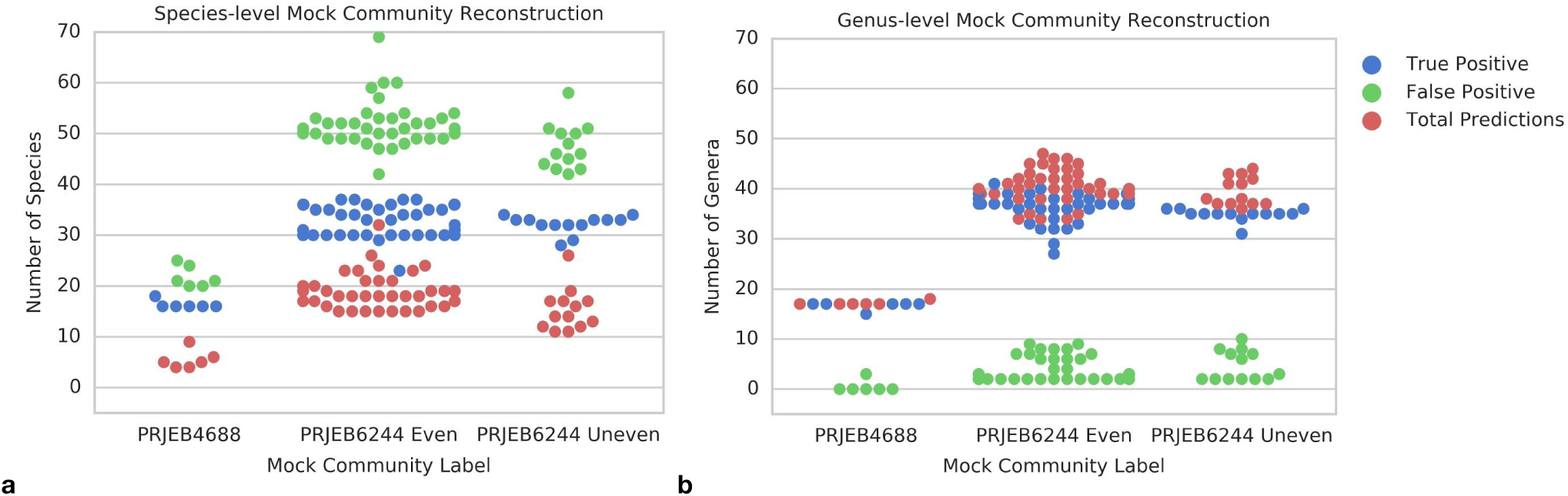
Mock community reconstructions using our deep learning approach. **(a)** The number of total predictions, true positives, and false positives contained in the species-level reconstructions calculated using model-based estimation with DNN_100_ (trained with 4% base-flipping noise) for 6 replicates of the 20-organism mock community from ENA study PRJB4688 and 51 runs of the 59-organism community from PRJEB6244. **(b)** The same as **(a)** for the genus-level reconstructions.

## 4 Discussion

We proposed a deep learning method that performs at or above state-of-the-art levels on a specific sequence-labeling problem: predicting species of origin for individual reads of 16S ribosomal DNA. We demonstrated this DNN is capable of near-optimal species classification, and moreover that these species-level predictions marginalize accurately to higher taxonomic ranks (Figure 2). When we examined the DNN’s individual predictions along a fixed reference sequence, the majority of confident, correct species-level predictions were achieved for queries covering parts of the 16S hypervariable regions (Appendix 5). As these hypervariable regions are well-known and exploited features of 16S sequences,^44,45^ the ability to learn the importance of these regions from training data is an important demonstration that 1) the DNN learned important biological structure of 16S, and 2) we might be able to discover unknown structure by inspecting the DNN’s behavior. We also validated that our approach works on experimental mock community data produced using typical Illumina sequencing protocols (Figure 5).

Our DNNs perform well compared to a diverse set of existing programs. They achieved read-level order classification accuracy comparable to Khawaldeh et al.^11^ despite modeling 22-fold more distinct orders (Figure 2b), and assigned genera to reads from novel species more accurately than naive Bayes approach based on the RDP Classifier^8^ (Figure 3). Though alignment baselines were slightly more competitive, our DNNs classified reads from held-out species more accurately for almost every read length we tried and were more robust to noise (Figure 3). This suggests our approach of injecting random base-flipping noise at train time allows the DNNs to learn a more robust noise model than the one implicitly incorporated by aligners like BLAST and BWA by allowing gaps and mismatches to be tolerated. Similarly, compared to Kraken2, our DNNs achieve PPVs which degrade less with increasing error rates and strike a better balance between sensitivity and precision overall (Figure 4). Therefore, we demonstrated that our deep learning approach is robust to high error rates where other popular approach struggle, an improvement which is particularly important considering newer technologies which produce more errorful reads, such as those being developed by Pacific Biosciences and Oxford Nanopore Technologies.^43^

Runtime performance is also a critical element of sequence-labelers; tools like Kraken2 are heavily optimized for speed. Our model architecture uses depthwise separable convolutions, which have been shown empirically to use parameters more efficiently than regular convolutions.^12-14^ The DNNs used in the current study consists of roughly 1.2 million floating-point operations per query and thus, though actual performance likely vary by application, common hardware accelerators for machine learning could allow them to process roughly 13 million (on a V100 GPU) to 156 million (on a Cloud TPU) reads per second.Moreover, our modeling choices make our DNNs well-suited for adaptation to mobile hardware, in contrast to approaches like alignment requiring access to large databases.

While our initial results are highly suggestive that downstream metagenomic analyses could benefit from complementing or replacing existing sequence-labeling substeps with our deep learning approach, more work is needed to fully understand its behavior. The current work does not explore performance on paired data, though it would be simple to extend our model to leverage this data by concatenating the reads. Additional adjustments for factors like unequal coverage in training data may be needed to properly calibrate the models’ predicted probability estimates (Appendix 5). More generally, it is possible that other deep learning architectures could further improve performance. When applying our approach to full metagenomic data, it might be necessary to incorporate some mechanism for detecting queries outside the training data’s domain, similar to the way Kraken2 attains higher PPV by leaving some queries unclassified. Finally, while our study on the effect of base-flips implies robustness of the method against sequencing errors and mutation rates, some of the higher error rates where our DNNs make the largest improvement may not be realistic for widely-used sequencing technologies like Illumina; follow-up analyses should specifically explore the behavior of our approach on reads produced by more errorful sequencers.

We emphasize that the model and methods proposed here are highly general, making this new approach an excellent candidate for complementing or replacing solutions to many large-scale bioinformatics problems which currently rely on string-matching or less flexible statistical modeling. Due to the improved robustness of our deep learning approach, impact may highest for applications where data is inherently noisy including analysis of bisulfite sequencing, viral DNA, immune-mapping, or ancient DNA. Overall, the current study acts as a first step towards our long-term goal of developing a general-purpose deep learning model that can successfully perform any task framed as the assignment of labels to short biological sequences.

## Acknowledgements

For their early input, which helped to frame the initial problem and understand potential applications, we thank Adam Roberts, Cinjon Resnick, and C. Rob Young. For their sharing of experimentalist perspectives on the problem, we thank Mauricio Carneiro, Vanessa Ridaura, and Roie Levy. This work was supported by internal funding.

