## Supplementary materials for "A deep learning approach to pattern recognition for short DNA sequences"

### Appendix 1: NCBI Data

| Phylum | References | Species | Genera | Families | Orders | Classes |
|---|---|---|---|---|---|---|
| Proteobacteria | 7,053 | 5,061 | 1,106 | 158 | 55 | 9 |
| Actinobacteria | 4,768 | 3,313 | 383 | 68 | 29 | 6 |
| Firmicutes | 3,814 | 2,531 | 499 | 56 | 13 | 7 |
| Bacteroidetes | 1,934 | 1,525 | 360 | 39 | 8 | 7 |
| Euryarchaeota | 834 | 450 | 100 | 28 | 13 | 8 |
| Tenericutes | 266 | 195 | 8 | 5 | 4 | 1 |
| Spirochaetes | 146 | 100 | 16 | 6 | 4 | 1 |
| Deinococcus-Thermus | 118 | 99 | 9 | 3 | 2 | 1 |
| Crenarchaeota | 113 | 61 | 27 | 8 | 5 | 1 |
| Cyanobacteria | 113 | 88 | 59 | 30 | 8 | 2 |
| Fusobacteria | 75 | 37 | 10 | 2 | 1 | 1 |
| Thermotogae | 70 | 48 | 13 | 5 | 4 | 1 |
| Verrucomicrobia | 57 | 53 | 22 | 7 | 4 | 3 |
| Acidobacteria | 45 | 41 | 19 | 5 | 5 | 4 |
| Planctomycetes | 44 | 31 | 23 | 5 | 3 | 2 |
| Chloroflexi | 44 | 35 | 26 | 15 | 12 | 8 |
| Aquificae | 43 | 32 | 14 | 4 | 2 | 1 |
| Synergistetes | 31 | 25 | 15 | 1 | 1 | 1 |
| Chlamydiae | 28 | 18 | 7 | 5 | 2 | 1 |
| Chlorobi | 21 | 16 | 5 | 1 | 1 | 1 |
| Deferribacteres | 15 | 11 | 7 | 1 | 1 | 1 |
| Thermodesulfobacteria | 14 | 12 | 5 | 1 | 1 | 1 |
| Nitrospirae | 11 | 10 | 3 | 1 | 1 | 1 |
| Fibrobacteres | 10 | 4 | 3 | 3 | 3 | 3 |
| Balneolaeota | 9 | 9 | 4 | 1 | 1 | 1 |
| Chrysiogenetes | 6 | 4 | 3 | 1 | 1 | 1 |
| Lentisphaerae | 6 | 5 | 3 | 3 | 3 | 2 |
| Dictyoglomi | 5 | 2 | 1 | 1 | 1 | 1 |
| Rhodothermaeota | 5 | 5 | 3 | 2 | 1 | 1 |
| Gemmatimonadetes | 5 | 4 | 3 | 2 | 2 | 2 |
| Ignavibacteriae | 4 | 2 | 2 | 2 | 1 | 1 |
| Armatimonadetes | 4 | 3 | 3 | 3 | 3 | 3 |
| Caldiserica | 3 | 1 | 1 | 1 | 1 | 1 |
| Calditrichaeota | 2 | 2 | 1 | 1 | 1 | 1 |
| Thaumarchaeota | 2 | 2 | 2 | 2 | 2 | 2 |
| Elusimicrobia | 2 | 1 | 1 | 1 | 1 | 1 |
| Kiritimatiellaeota | 1 | 1 | 1 | 1 | 1 | 1 |
| Nitrospinae | 1 | 1 | 1 | 1 | 1 | 1 |

**Extended Data Table 1: NCBI dataset breakdown by phylum.** The distribution of reference sequences and species, genus, family, order, and class labels across the 38 different phyla represented in our NCBI dataset.

Our NCBI dataset is based on 19,851 16S ribosomal RNA sequences (18,902 bacterial and 949 archaeal) which have an average length of 1,454.13 base pairs, with individual sequences varying from 302 to 3,600 base pairs. During the labelling process, we excluded 129 reference sequences whose reported taxonomic labels in the NCBI Taxonomy Browser violated the tree structure of the overall taxonomy. The resulting dataset includes 13,838 distinct species which are distributed across 38 phyla according to Extended Data Table 1. Subsequences from these 19,722 references comprise our synthetic $NCBI_L$ read sets. The total number of synthetic reads contained in each of our $NCBI_L$ datasets is determined by the length *L*: $NCBI_L$ contains 28,219,784, 27,726,734, 26,740,634, 25,754,534, and 24,768,434 reads for lengths 25, 50, 100, 150, and 200, respectively.

### Appendix 2: Model Architecture and Implementation

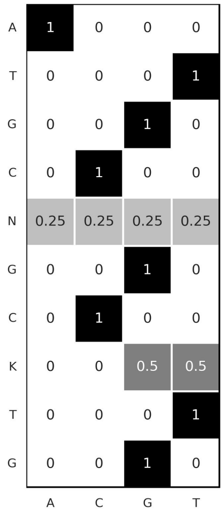


**Extended Data Figure 1: Example input encoding.** Input encoding for a sample 10 base pair sequence demonstrating how both canonical bases and IUPAC ambiguity codes are encoded using four-dimensional probability distributions.

For our models, we used the input representation in Extended Data Figure 1. Each read is a short sequence of canonical nitrogenous bases (A, C, T, G) and IUPAC ambiguity codes (K, M, R, Y, S, W, B, V, H, D, X, N). We one-hot encoded each canonical base as a four-dimensional vector and resolved each ambiguity code to the appropriate probability distribution over these four bases. Note that this approach to input encoding does not make use of any quality scores; it would be straightforward to extend our approach to include this information, for example by using an extra input channel.

A key feature of our proposed model architecture is its use of depthwise separable convolutions. Initially studied by Sifre & Mallat (2013), depthwise separable convolutions separate the task of learning spatial features from that of integrating information across channels by decomposing a typical convolution into two sequential operations: a spatial convolution applied independently over each input channel followed by a pointwise convolution across channels. We use of 1D depthwise separable convolutions, formalized as follows given input *x* with *C* channels and a filter of width *F*:

$$PointwiseConv(W,x)_{(i)} = \sum_{c}^{C} W_c \cdot x_{(i,c)}$$

$$DepthwiseConv(W,x)_{(i)} = \sum_{f}^{F} W_f \circ x_{(i+f)}$$

$$SeparableConv(W_p, W_d, x)_{(i)} = PointwiseConv_{(i)}(W_p, DepthwiseConv_{(i)}(W_d, x))$$

where *W* denotes a weight matrix and $\circ$ is element-wise multiplication.

After each convolutional and fully-connected layer, we use the following version of leaky rectified-linear activation (Maas, Hannun, & Ng, 2013; Xu, et al., 2015) after every convolutional and fully-connected layer:

$$LReLU(x)_i = max(x_i,\ ax_i) = \{x_i \ if\ x_i \geq 0,\ ax_i \ if\ x_i < 0$$

where the slope $a \in (0, 1)$ for each model is as in Extended Data Table 2.

| Name | Training Read Length | Spatial Conv Widths | Pointwise Conv Depths | Number FC Layers | Number FC Units | lReLU Slope | Learning Rate | Decay Rate | Keep Prob | Weight Init Scale |
|---|---|---|---|---|---|---|---|---|---|---|
| $DNN_{25}$ | 25 | 13, 9, 9 | 34, 48, 37 | 3 | 2,969 | $1.1619e^{-2}$ | $5.2225e^{-4}$ | $5.0277e^{-2}$ | 87.107% | 1.6181 |
| $DNN_{50}$ | 50 | 13, 35, 13 | 51, 151, 114 | 2 | 2,919 | $1.1699e^{-2}$ | $7.4034e^{-4}$ | $7.8038e^{-2}$ | 89.161% | 2.2217 |
| $DNN_{100}$ | 100 | 5, 9, 13 | 84, 58, 180 | 2 | 2,828 | $1.2538e^{-2}$ | $4.6969e^{-4}$ | $6.5505e^{-2}$ | 94.018% | 1.1841 |
| $DNN_{150}$ | 150 | 5, 9, 21 | 59, 221, 119 | 3 | 2,908 | $5.7478e^{-3}$ | $7.5135e^{-5}$ | $9.1889e^{-2}$ | 88.834% | 2.4636 |
| $DNN_{200}$ | 200 | 9, 5, 21 | 197, 116, 119 | 2 | 2,733 | $1.1491e^{-2}$ | $6.7080e^{-4}$ | $6.4534e^{-2}$ | 91.967% | 0.5878 |

**Extended Data Table 2: Selected neural network hyperparameters.** The best deep neural network (DNN) model hyperparameters identified for each read length $L$={25, 50, 100, 150, 200}.

During training, we initialized each layer's parameters according to a truncated random normal distribution with standard deviation $S/\sqrt{N}$ , where $S$ is the weight initialization scale in Extended Data Table 2 and $N$ is the number of inputs to the layer. On each parameter update, we clipped the gradients to have norm at most 20.

### Appendix 3: Data Splits and Model Selection

For model selection, we split our $NCBI_L$ datasets into three smaller subsets: NCBI-$0_L$, NCBI-$1_L$, and NCBI-$2_L$. We constructed NCBI-$0_L$ by first taking a random sample of 90% of the species in each genus (selecting at least one species per genus), then sampling 90% of the reads for each selected species. The remaining 10% of the reads for these species form NCBI-$1_L$, and NCBI-$2_L$ contains all the reads for the 10% of species excluded from NCBI-$0_L$. As an example, Extended Data Table 3 enumerates the contents of our NCBI-$0_{100}$, NCBI-$1_{100}$, and NCBI-$2_{100}$ subsets.

| NCBI-$0_{100}$ | | NCBI-$1_{100}$ | | NCBI-$2_{100}$ | |
|---|---|---|---|---|---|
| 90% of reads from 90% of species per genus | | Remaining 10% of reads from species in NCBI0 | | 100% of reads from remaining 10% of species per genus | |
| Reads | 21,899,715 | Reads | 2,431,551 | Reads | 2,409,368 |
| Superkingdoms | 2 | Superkingdoms | 2 | Superkingdoms | 2 |
| Phyla | 38 | Phyla | 38 | Phyla | 23 |
| Classes | 91 | Classes | 91 | Classes | 48 |
| Orders | 202 | Orders | 202 | Orders | 110 |
| Families | 479 | Families | 479 | Families | 227 |
| Genera | 2,768 | Genera | 2,768 | Genera | 577 |
| Species | 12,609 | Species | 12,609 | Species | 1,229 |

**Extended Data Table 3: NCBI subset contents for 100 base pair data.** The contents of each subset of our $NCBI_{100}$ dataset in terms of the total number of reads and the number of distinct labels at each taxonomic rank.

For $L$ = {25, 50, 100, 150, 200} we selected a model $DNN_L$ by training on noiseless reads from NCBI-$0_L$ and performing a hyperparameter search to maximize read-level accuracy on a validation set comprised of the reads in NCBI-$1_L$ and NCBI-$2_L$ with base-flipping noise injected at a rate of 1%. Because reads in NCBI-$2_L$ are from species held out during training, we measured read-level accuracy on this validation set as follows:

1. If the current example arose from the reference sequence of a species represented in NCBI-$0_L$, the prediction by $DNN_L$ is correct if the model assigns the most probability mass to the true species label.
2. Otherwise, if the example arose from the reference sequence of a held-out species, the model's prediction is correct if the true genus label receives the most probability mass when the model's output is marginalized to the genus-level distribution.

We used Google Vizier (Golovin, et al., 2017) with the default search algorithm to explore the hyperparameter space and optimize the objective computed in this manner. The hyperparameters in Extended Data Table 2 are the best ones discovered by this search.

### Appendix 4: Pooling Studies

In the current work, we explored one straightforward approach for handling the variable read lengths produced by next-generation sequencing technologies: we enabled running a given model on any query at least as long as the fixed input dimension of the fully-connected layers by tiling the fully-connected layers and adding a pooling layer between the last fully-connected layer and the softmax output layer. We determined which type of pooling works best on our species classification problem by training several models on our NCBI-$0_{100}$ data with the effective width of the fully-connected layers set to 80 base pairs to trigger modeling tiling and pooling. For both average and max pooling, we fixed the number of depthwise separable convolutional layers to 1 and performed random search over the remaining hyperparameters. We trained each model for 200,000-400,000 training iterations and then evaluated on a validation set as described in Appendix 3.

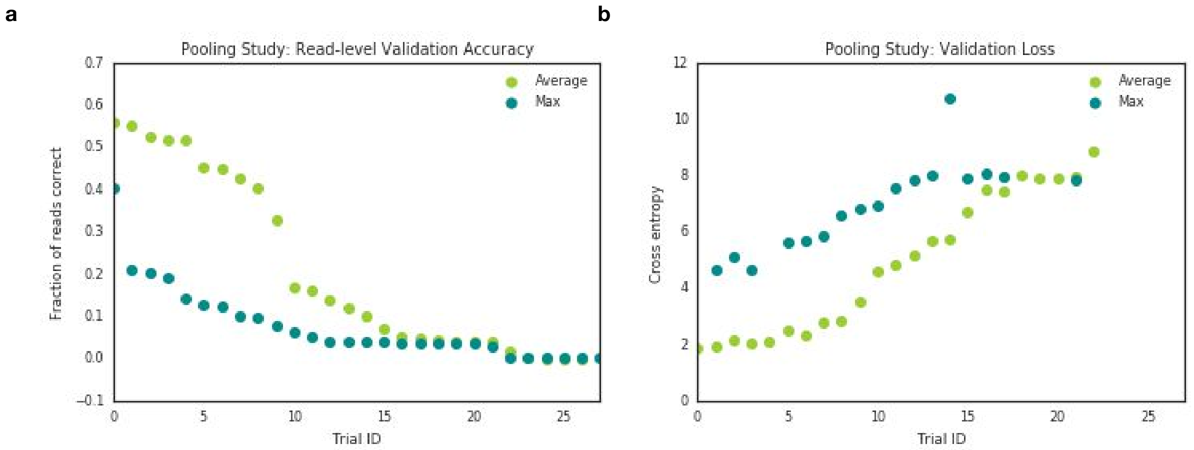


**Extended Data Figure 2: Differences in accuracy and cross entropy loss for average and max pooling.** Performance comparison for average and max pooling trials for both **(a)** read-level accuracy and **(b)** cross entropy loss on validation data, where trial IDs (x-axis) are assigned according to descending validation accuracy.

In total, we trained 40 models with max pooling and 27 with average pooling. Despite the skew in number of attempted trials, we found models with average pooling to perform significantly better than those with max pooling; the models with average pooling attained higher accuracies and lower loses at both train and evaluation time (Extended Data Figure 2). 27 of the models with max pooling layers either failed to converge or never outperformed random guessing, compared to only 9 such models with average pooling. The head-to-head comparison of the highest achieved read-level accuracies on the validation set further reveals the extent of this performance differential: the best average pooling model outperforms the best max pooling model by more than 34.6%. This pooling exploration established a sound method of constructing relatively flexible models for read-level species classification of 16S sequencing data. Based on these results, all models presented in the current study use average pooling.

### Appendix 5: Hypervariable Regions

While read-level classification rates are a key measure of the deep learning models' performance on our taxonomic classification benchmarking problem, they obscure the probabilistic nature of their predictions. To investigate whether these probability assignments themselves have any interesting properties, we compared the probability weights assigned by $DNN_{100}$ across the length of a fixed reference sequence. Extended Data Figure 3a shows that $DNN_{100}$ made no within-genus mistakes on synthetic reads from the *Salmonella enterica* reference sequence, whereas in Extended Data Figure 3b within-genus mistakes were common in regions of the reference sequence where $DNN_{100}$ assigned low probability weights to the true *Salmonella bongori* label. In both of these cases we found that the neural network made its most confident predictions on 100 base pair synthetic reads containing portions of the hypervariable regions (as identified using analysis by Chakravorty et al. (2007) and *E. coli* coordinates from Brosius et al. (1978)). Thus, although the model did not learn to perfectly recapitulate every hypervariable region (for example, it made mistaken species assignments on reads containing portions of the V5 region in Extended Data Figure 3b), it nonetheless appears to have learned to make its most accurate, confident predictions within the hypervariable regions. Indeed, the lack of overly confident predictions on reads from purely conserved regions of the reference sequences further suggests the model has learned from reasonable signal in the data rather than overfitting to artifacts present in our particular set of training reference sequences.

However, when we repeated this analysis on a reference sequence for *Streptomyces libani*, one of the 681 distinct species from the most prevalent genus in the training set, we found that the model's predicted probabilities followed an entirely different trend (Extended Data Figure 3c and d). Unlike in the *Salmonella* cases, probabilities assigned to the correct species label are low across the entire reference and within-genus mistakes dominated. Extended Data Figure 3d, on the other hand, shows that the model's genus-level predictions were both confident and accurate on the same synthetic reads, with $DNN_{100}$ making no genus mistakes and assigning at least 0.8 probability to the true *Streptomyces* label everywhere except for a small region between V2 and V3. This suggests that predicted probability mass is divided amongst multiple closely-related species which cannot be disambiguated due to an insufficient proportion of distinctive 100 base pair segments within their reference sequences, so that the particular region a given synthetic read was pulled from does not appear to matter. This issue is likely caused or exacerbated by the fact that no adjustments were made for differential dataset coverage.

Examining the DNN's confidence in its read-mapping assignments along the length of 16S references provides some initial evidence that these models achieve their good performance by learning salient read-level information. Extended Data Figure 3 shows that, in general, the vast majority of queries which yield confident, correct species-level predictions tend to cover parts of the hypervariable regions. However, it also exposes the impact that differential coverage in the training set may have on individual predictions: the model made many within-genus mistakes for the less prevalent *Salmonella bongori* (2 references) but none for *Salmonella enterica* (11 references), a trend which was even more apparent for *Streptomyces libani*, one of the species from the most prevalent genus in our dataset. A more robust training or inference scheme which properly adjusts for skewed coverage might improve the quality of these individual predictions and allow the model's predicted probability estimates to be leveraged to give calibrated confidence estimates for label assignments.

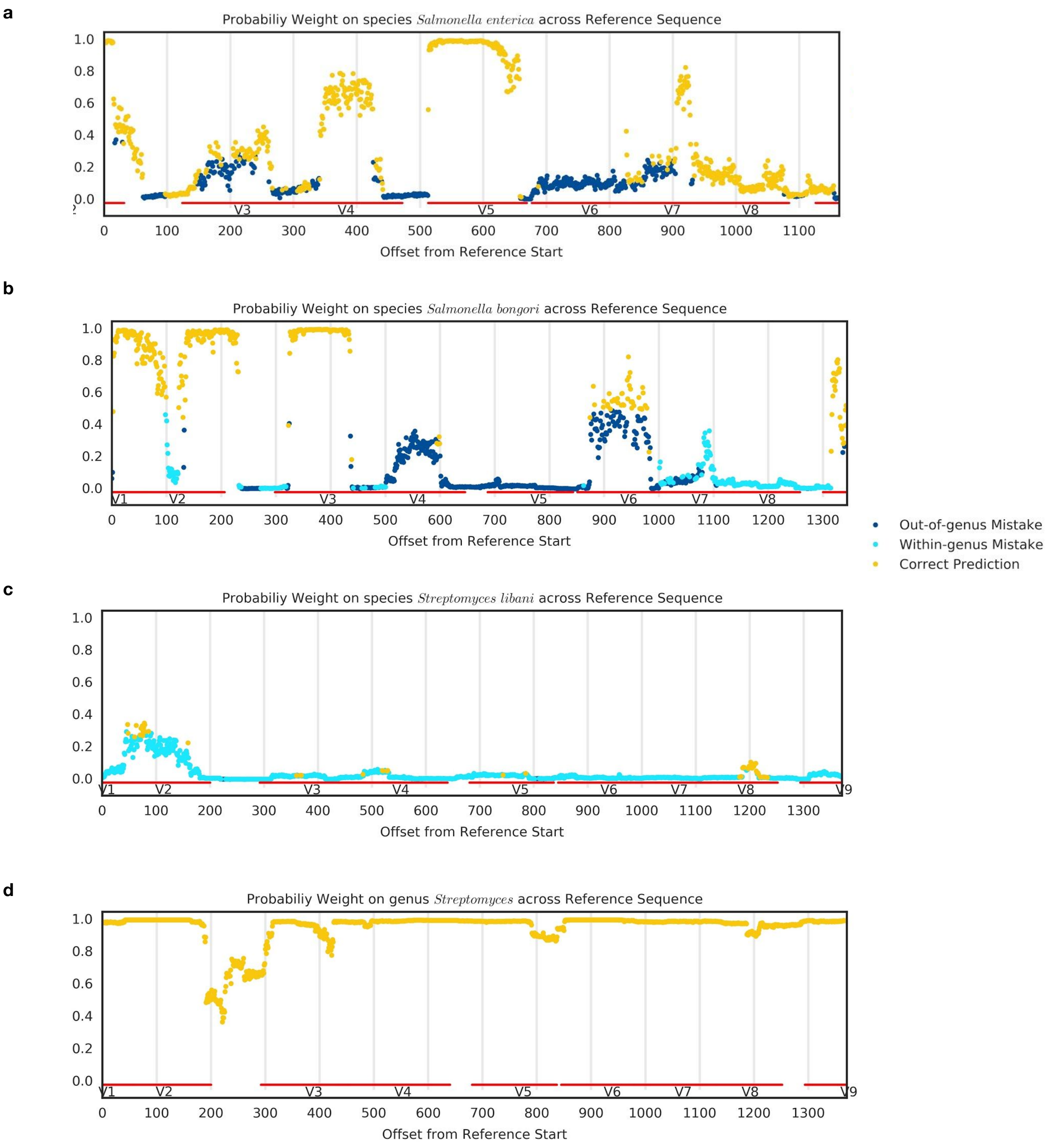


**Extended Data Figure 3: Variations in confidence of deep learning approach along fixed 16S reference sequences.** Probability weight assigned to the correct species label by $DNN_{100}$ for every 100 base pair subsequence of **(a)** *Salmonella enterica* (RefSeq ID NR_116126.1), **(b)** *Salmonella bongori* (NR_116124.1), and **(c)** *Streptomyces libani* (NR_042301.1) reference sequences. Offset from the beginning of the reference to the start of the subsequence is specified on the x-axis, and color represents whether $DNN_{100}$'s most confident prediction is the correct label (yellow), another species label in the correct genus (cyan), or a species outside the genus (blue). **(d)** The genus-level probability weights assigned by $DNN_{100}$ to the Streptomyces label for the same reference sequence as **(c)**.

### Appendix 6: Baseline Methods

We computed alignment baselines based on BLAST and BWA mappings against the original reference sequences. Taking *T* to be a fixed set of short reads, we computed the BWA baseline as follows:

1. Set $accuracy = 0$ and for each read $x \in T$:
    a. Use BWA to assign a set of mappings *A* and primary mapping $a^*$ to the read *x.* For each alignment mapping *a*, let $a_{ref}$ be the reference sequence involved in the alignment and $a_{ed}$ be the corresponding edit distance score.
    b. If *A* is non-empty:
        i. Take $A^* = \{a \in A \mid a_{ed} \leq a^*_{ed}\}$
        ii. $accuracy \pm \frac{C}{|A^*|}$ where *C* is the number of times the true label for read *x* appears in the set of ground-truth labels for $\{a_{ref} \mid a \in A^*\}$
2. Take $\frac{accuracy}{|T|}$ as the final accuracy rate

The BLAST baseline accuracy replaces the comparison in 1b with one checking for bit scores that are at least as large as the bit score of the best mapping.

Our naive Bayes classifier was based on the RDP Classifier, and as such, we used the 8-mer representation and computed prior probabilities and genus-specific conditional probabilities as described in Wang et al. (2007) We additionally devised a method of handling IUPAC ambiguity codes in the 8-mer vector representation by assigning each possible DNA 8-mer encoded by the IUPAC code a fractional presence. For example, the 9 base pair sequence 'AAAAAAAAN' was transformed into a vector with four non-zero entries: AAAAAAAA with weight 1.0, and AAAAAAAC, AAAAAAAG, and AAAAAAAT with weights 0.25. We found that incorporating these ambiguous bases improved accuracy for short or noisy reads.

For our Kraken2 baseline, we generated *names.dmp* and *nodes.dmp* files corresponding to the taxonomy of organisms in our NCBI data, and a corresponding FASTA file marked with Kraken2 *taxid* markers for the 16S reference sequences corresponding to the $NCBI_L$-0 training sets. We then constructed an appropriate Kraken2 database using the following commands:

```
$ kraken2-build --db kraken2_database --add-to-library kraken.fasta --no-masking
$ kraken2-build --db kraken2_database --build --kmer-len ${kmer_len}
--minimizer-len ${minimizer_len} --minimizer-spaces ${minimizer_spaces}
```

We set *kmer_len*, *minimizer_len*, and *minimizer_spaces* to their default values of (35, 31, 6) for running on 50, 100, and 200 base pair reads and to (15, 15, 3) for running on 25 base pair reads since in this setting the default values are too large.

### Appendix 7: Mock Community Data

The sequencing data from study PRJEB4688 comes from a community developed by the Human Microbiome Project (Huttenhower et al., 2012) to contain equal concentrations of the following 20 bacterial species: *Acinetobacter baumannii str. 5377*, *Actinomyces odontolyticus str. 1A.21*, *Bacillus cereus str. NRS 248*, *Bacteroides vulgatus str. NCTC 11154*, *Clostridium beijerinckii str. NCIMB 8052*, *Deinococcus radiodurans str. R1, (smooth)*, *Enterococcus faecalis str. OG1RF*, *Escherichia coli str. K12 substr. MG1655*, *Helicobacter pylori str. 26695*, *Lactobacillus gasseri str. 63 AM*, *Listeria monocytogenes str. EGDe*, *Neisseria meningitidis str. MC58*, *Propionibacterium acnes str. KPA171202*, *Pseudomonas aeruginosa str. PAO1-LAC*, *Rhodobacter sphaeroides str. ATH 2.4.1*, *Staphylococcus aureus TCH1516*, *Staphylococcus epidermidis FDA str. PCI 1200*, *Streptococcus agalactiae str. 2603 V/R*, *Streptococcus mutans str. UA159*, and *Streptococcus pneumoniae str. TIGR4*. There are six replicates in total: three single-ended and three paired-end replicates. The paired-end replicates, ERR619081-3, contain 481,364, 426,086, and 180,252 unpaired reads, respectively, all of length 251 base pairs. The single-ended replicates, ERR348713-5, contain reads of variable lengths ranging from 225 to 384 base pairs distributed according to Extended Data Table 4.

| ERR348713 | | ERR348714 | | ERR348715 | |
|---|---|---|---|---|---|
| **Read Length** | **Frequency** | **Read Length** | **Frequency** | **Read Length** | **Frequency** |
| 248 | 4 | 225 | 1 | 370 | 689 |
| 249 | 1 | 246 | 1 | 371 | 11,697 |
| 250 | 15 | 248 | 3 | 372 | 277 |
| 251 | 106 | 249 | 1 | 373 | 2,955 |
| 252 | 33,558 | 250 | 16 | 374 | 31,233 |
| 253 | 156,843 | 251 | 100 | 375 | 38,529 |
| 254 | 22,512 | 252 | 36,868 | 376 | 4,711 |
| 255 | 4 | 253 | 177,126 | 377 | 33 |
| | | 254 | 26,554 | 384 | 2 |
| | | 255 | 11 | | |
| | | 259 | 1 | | |
| all | 213,043 | all | 240,682 | all | 90,126 |

**Extended Data Table 4: Read length distribution for single-ended mock community replicates.** Distribution of read lengths for each single-ended replicate from ENA study PRJEB4688.

The mock community sequenced in study PRJEB6244, contains 59 distinct organisms. The even version of the community contains an equal number of molecules per strain, but there is also an uneven version for which strain amounts are log-normally distributed within each phylum. Extended Data Table 5 gives the specific strains and uneven concentrations according to previous publications (D'Amore et al. 2016; Schirmer et al., 2015). Adjusting for updated taxonomic assignments for some organisms, these 59 strains are covered by the following 56 species labels in NCBI: *Acidobacterium capsulatum*, *Akkermansia muciniphila*, *Bacteroides thetaiotamicron*, *Bacteroides vulgatus*, *Bordetella bronchiseptica*, *Caldicellulosiruptor bescii*, *Caldicellulosiruptor saccharolyticus*, *Chlorobaculum tepidum*, *Chlorobium limicola*, *Chlorobium phaeobacteroides*, *Chlorobium phaeovibrioides*, *Chloroflexus aurantiacus*, *Deinococcus radiodurans*, *Desulfovibrio desulfuricans*, *Desulfovibrio piger*, *Dickeya dadantii*, *Dictyoglomus turgidum*, *Enterococcus faecalis*, *Fusobacterium nucleatum*, *Gemmatimonas aurantiaca*, *Herpetosiphon aurantiacus*, *Hydrogenobaculum sp.*, *Leptothrix cholodnii*, *Nitrosomonas europaea*, *Nostoc sp.*, *Paraburkholderia xenovorans*, *Pelodictyon phaeoclathratiforme*, *Persephonella marina*, *Porphyromonas gingivalis*, *Rhodopirellula baltica*, *Rhodospirillum rubrum*, *Ruegeria pomeroyi*, *Ruminiclostridium thermocellum*, *Salinispora arenicola*, *Salinispora tropica*, *Shewanella baltica*, *Sulfitobacter sp.*, *Sulfurihydrogenibium sp.*, *Sulfurihydrogenibium yellowstonense*, *Thermoanaerobacter pseudethanolicus*, *Thermotoga neapolitana*, *Thermotoga petrophila*, *Thermotoga sp.*, *Thermus thermophilus*, *Treponema denticola*, *Treponema vincentii*, *Zymomonas mobilis*, *Archaeoglobus fulgidus*, *Ignicoccus hospitalis*, *Methanocaldococcus jannaschii*, *Methanococcus maripaludis*, *Nanoarchaeum equitans*, *Pyrobaculum aerophilum*, *Pyrobaculum calidifontis*, *Pyrococcus horikoshii*, and *Sulfolobus tokodaii*.

At the genus-level, these organisms are covered by 45 labels: *Acidobacterium*, *Akkermansia*, *Bacteroides*, *Bordetella*, *Caldicellulosiruptor*, *Chlorobaculum*, *Chlorobium*, *Chloroflexus*, *Deinococcus*, *Desulfovibrio*, *Dictyoglomus*, *Dickeya*, *Enterococcus*, *Fusobacterium*, *Gemmatimonas*, *Herpetosiphon*, *Hydrogenobaculum*, *Leptothrix*, *Nitrosomonas*, *Nostoc*, *Paraburkholderia*, *Pelodictyon*, *Persephonella*, *Porphyromonas*, *Rhodopirellula*, *Rhodospirillum*, *Ruegeria*, *Ruminiclostridium*, *Salinispora*, *Shewanella*, *Sulfitobacter*, *Sulfurihydrogenibium*, *Thermoanaerobacter*, *Thermotoga*, *Thermus*, *Treponema*, *Zymomonas*, *Archaeoglobus*, *Ignicoccus*, *Methanocaldococcus*, *Methanococcus*, *Nanoarchaeum*, *Pyrobaculum*, *Pyrococcus*, and *Sulfolobus*.

| Name | Domain | Proportion | Name | Domain | Proportion |
|---|---|---|---|---|---|
| *Acidobacterium capsulatum ATCC 51196* | Bacteria | 8.1% | *Rhodopirellula baltica SH 1* | Bacteria | 1.0% |
| *Akkermansia muciniphila ATCC BAA-835* | Bacteria | 0.9% | *Rhodospirillum rubrum ATCC 11170* | Bacteria | 1.2% |
| *Anaerocellum thermophilum Z-1320, DSM 6725* | Bacteria | 1.2% | *Ruegeria pomeroyi DSS-3* | Bacteria | 0.6% |
| *Bacteroides thetaiotaomicron VPI-5482* | Bacteria | 0.2% | *Salinispora arenicola CNS-205* | Bacteria | 0.5% |
| *Bacteroides vulgatus ATCC 8482* | Bacteria | 0.9% | *Salinispora tropica CNB-440* | Bacteria | 1.6% |
| *Bordetella bronchiseptica RB50* | Bacteria | 9.2% | *Shewanella baltica OS185* | Bacteria | 3.1% |
| *Burkholderia xenovorans LB400* | Bacteria | 2.6% | *Shewanella baltica OS223* | Bacteria | 1.4% |
| *Caldicellulosiruptor saccharolyticus DSM 8903* | Bacteria | 2.0% | *Sulfitobacter sp. EE-36* | Bacteria | 2.0% |
| *Chlorobaculum tepidum TLS* | Bacteria | 0.5% | *Sulfitobacter sp. NAS-14.1* | Bacteria | 4.3% |
| *Chlorobium limicola DSM 245* | Bacteria | 0.4% | *Sulfurihydrogenibium sp. YO3AOP1* | Bacteria | 1.6% |
| *Chlorobium phaeobacteroides DSM 266* | Bacteria | 1.9% | *Sulfurihydrogenibium yellowstonense SS-5* | Bacteria | 2.6% |
| *Chlorobium phaeovibrioides DSM 265* | Bacteria | 0.3% | *Thermoanaerobacter pseudethanolicus ATCC 33223* | Bacteria | 0.8% |
| *Chloroflexus aurantiacus J-10-fl* | Bacteria | 0.9% | *Thermotoga neapolitana DSM 4359* | Bacteria | 0.7% |
| *Clostridium thermocellum ATCC 27405* | Bacteria | 0.6% | *Thermotoga petrophila RKU-1* | Bacteria | 1.0% |
| *Deinococcus radiodurans R1* | Bacteria | 1.7% | *Thermotoga sp. RQ2* | Bacteria | 3.4% |
| *Desulfovibrio desulfuricans ATCC 27774* | Bacteria | 1.4% | *Thermus thermophilus HB8* | Bacteria | 0.5% |
| *Desulfovibrio piger ATCC 29098* | Bacteria | 3.1% | *Treponema denticola ATCC 35405* | Bacteria | 0.2% |
| *Dictyoglomus turgidum DSM 6724* | Bacteria | 3.5% | *Treponema vincentii I* | Bacteria | 0.2% |
| *Erwinia chrysanthemi* | Bacteria | 0.3% | *Zymomonas mobilis mobilis ZM4* | Bacteria | 0.8% |
| *Enterococcus faecalis V583* | Bacteria | 4.3% | *Archaeoglobus fulgidus DSM 4304* | Archaea | 0.3% |
| *Fusobacterium nucleatum ATCC 25586* | Bacteria | 0.3% | *Ignicoccus hospitalis KIN4/I* | Archaea | 1.2% |
| *Gemmatimonas aurantiaca T-27T* | Bacteria | 0.7% | *Methanocaldococcus jannaschii DSM 2661* | Archaea | 0.9% |
| *Herpetosiphon aurantiacus ATCC 23779* | Bacteria | 1.8% | *Methanococcus maripaludis C5* | Archaea | 0.4% |
| *Hydrogenobaculum sp. Y04AAS1* | Bacteria | 1.1% | *Methanococcus maripaludis S2* | Archaea | 0.5% |
| *Leptothrix cholodnii SP-6* | Bacteria | 1.8% | *Nanoarchaeum equitans Kin4-M* | Archaea | 1.0% |
| *Nitrosomonas europaea ATCC 19718* | Bacteria | 4.3% | *Pyrobaculum aerophilum IM2* | Archaea | 0.5% |
| *Nostoc sp. PCC 7120* | Bacteria | 2.7% | *Pyrobaculum calidifontis JCM 11548* | Archaea | 2.6% |
| *Pelodictyon phaeoclathratiforme BU-1* | Bacteria | 0.1% | *Pyrococcus horikoshii OT3* | Archaea | 1.9% |
| *Persephonella marina EX-H1* | Bacteria | 5.5% | *Sulfolobus tokodaii 7(S311)* | Archaea | 0.7% |
| *Porphyromonas gingivalis ATCC 33277* | Bacteria | 0.2% | | | |

**Extended Data Table 5: True contents of 59-organism mock community.** List of 10 Archaea and 59 bacterial strains present in the mock community from ENA study PRJEB6244.

Some of the labels for this 59-organism mock community are missing from our NCBI dataset, namely the genus label *Nanoarchaeum* and the eight species labels *Hydrogenobaculum sp.*, *Leptothrix cholodnii*, *Nostoc sp.*, *Sulfitobacter sp.*, *Sulfurihydrogenibium sp.*, *Thermotoga sp.*, *Treponema vincentii*, and *Nanoarchaeum equitans*. For our analyses, we used all of the mock community sequencing runs included in study PRJEB6244 which amounts to the 51 runs listed in Extended Data Table 6. All of the reads contained in the data files for each of these runs are 250 base pairs, but the total number of reads per run varies from 3,290 to 3,506,882.

| Run Accession | Number of Reads (unpaired) | Community Type | Run Accession | Number of Reads (unpaired) | Community Type |
|---|---|---|---|---|---|
| ERR777676 | 413,164 | Even | ERR777718 | 28,704 | Uneven |
| ERR777677 | 77,748 | Even | ERR777719 | 23,526 | Uneven |
| ERR777678 | 331,464 | Even | ERR777720 | 49,458 | Uneven |
| ERR777695 | 1,187,736 | Even | ERR777721 | 62,430 | Uneven |
| ERR777696 | 3,506,882 | Even | ERR777722 | 44,192 | Uneven |
| ERR777697 | 3,268,324 | Even | ERR777726 | 711,606 | Even |
| ERR777698 | 2,185,152 | Even | ERR777727 | 667,110 | Even |
| ERR777699 | 26,282 | Even | ERR777728 | 612,894 | Even |
| ERR777700 | 5,730 | Even | ERR777729 | 1,316,194 | Uneven |
| ERR777701 | 4,052 | Even | ERR777730 | 43,378 | Even |
| ERR777702 | 736,790 | Even | ERR777731 | 48,706 | Even |
| ERR777703 | 184,040 | Even | ERR777732 | 3,299,128 | Even |
| ERR777704 | 584,670 | Even | ERR777733 | 1,910,258 | Even |
| ERR777705 | 2,810,390 | Even | ERR777734 | 13,224 | Even |
| ERR777706 | 2,263,688 | Even | ERR777735 | 5,272 | Even |
| ERR777707 | 553,630 | Uneven | ERR777736 | 815,732 | Even |
| ERR777708 | 2,280,446 | Uneven | ERR777737 | 488,286 | Even |
| ERR777709 | 2,017,580 | Uneven | ERR777738 | 403,586 | Even |
| ERR777710 | 2,162,570 | Even | ERR777739 | 969,458 | Even |
| ERR777711 | 3,120,284 | Uneven | ERR777740 | 62,898 | Uneven |
| ERR777712 | 3,510 | Even | ERR777741 | 524,110 | Uneven |
| ERR777713 | 15,158 | Even | ERR777742 | 464,972 | Uneven |
| ERR777714 | 3,290 | Even | ERR777746 | 24,690 | Even |
| ERR777715 | 70,706 | Even | ERR777747 | 62,594 | Even |
| ERR777716 | 91,702 | Even | ERR777748 | 91,438 | Even |
| ERR777717 | 100,696 | Even | | | |

**Extended Data Table 6: Community type by run accession.** List of ENA accessions for mock community sequencing runs from study PRJEB6244 along with their corresponding community type (even or uneven) and read count.
